# Single-molecule DNA unzipping reveals asymmetric modulation of a transcription factor by its binding site sequence and context

**DOI:** 10.1101/188458

**Authors:** Sergei Rudnizky, Hadeel Khamis, Omri Malik, Allison Squires, Amit Meller, Philippa Melamed, Ariel Kaplan

## Abstract

Most functional transcription factor (TF) binding sites deviate from their “consensus” recognition motif, although their sites and flanking sequences are often conserved across species. Here, we used single-molecule DNA unzipping with optical tweezers to study how Egr-1, a TF harbouring 3 zinc fingers (ZF1,ZF2 and ZF3), is modulated by the sequence and context of its functional sites in the *Lhb* gene promoter. We find that both the core 9 base pairs bound to Egr-1 in each of the sites, and the base pairs flanking them, modulate the affinity and structure of the protein-DNA complex. The effect of the flanking sequences is asymmetric, with a stronger effect for the sequence flanking ZF3. Characterization of the dissociation time of Egr-1 revealed that a local, mechanical perturbation of the interactions of ZF3 destabilizes the complex more effectively than a perturbation of the ZF1 interactions. Our results reveal a novel role for ZF3 in the interaction of Egr-1 with other proteins and the DNA, providing insight on the regulation of *Lhb* and other genes by Egr-1. Moreover, our findings reveal the potential of small changes in DNA sequence to alter transcriptional regulation, and may shed light on the organization of regulatory elements at promoters.

## INTRODUCTION

Binding of Transcription Factors (TFs) to regulatory elements at the promoters and enhancers of genes is a central event in the regulation of transcription, and lies at the core of the cell’s response to environmental stimuli. As such, a detailed understanding of the interactions of TFs with DNA is crucial, not only at “large” scales, e.g. identifying TF binding sites throughout genomes(1–3), but also at the “small” scale, by developing a molecular understanding of the different factors affecting the structure and dynamics of the TF-DNA complex, and how they affect transcriptional regulation. Recent studies have highlighted the existence of multiple layers of complexity in the modulation of a TF binding to DNA, revealing the effects of the structure and flexibility of the binding site(4, 5), the kinetic competition of TFs with nucleosomes(6–8), cooperativity between TFs(9–13), the presence of neighboring TF sites(14, 15), and DNA methylation(16). However, the diversity of mechanisms by which TFs select binding sites *in vivo*and alter gene expression remain unclear, calling for a mechanistic understanding of all these factors to understand TF function.

According to numerous experimental observations, proteins find their specific binding sites on DNA very quickly and efficiently, with search times that are, in some cases, shorter than the ones estimated using 3D diffusion models(14, 15, 17, 18). This phenomena, known as “facilitated diffusion”, has stimulated the development of theoretical frameworks of the search process (reviewed in (15)). In particular, the model by Berg, Winter and von Hippel(19–21) postulates that proteins are able to find their targets by coupling three-dimensional diffusion in the solution, with one-dimensional diffusion while bound nonspecifically to DNA. Notably, although this theoretical approach is able to explain qualitative features of the facilitated diffusion, the measured diffusion constants and partitioning of time between the solution and the DNA call into question the utilization of this model to describe real biological system, and indicate that this simple model is not enough to rationalize the fast search dynamics(15). However, the sliding of proteins on DNA has been experimentally observed in single molecule experiments(14, 22–28), indicating that one-dimensional diffusion on the DNA is an integral part of the search process. This highlights an additional intriguing aspect of the search process: To be able to scan the DNA at high speeds, the protein binding potential must be a smooth function of the position on the DNA, with variations no larger that 1-2 k_b_T as a function of sequence. However, the required stability of the protein–DNA complex when bound at the specific site requires a tight protein-DNA interaction, i.e. a binding energy significantly higher than k_b_T, which would slow down the search process. This is commonly known as the “speed-stability” paradox(14, 29–31). A proposed solutions to this paradox postulates the existence of two different conformational states (and therefore different binding energies): one while the protein diffuses along the DNA, and another one when it probes the DNA for a target sequence(14, 29, 32, 33). However, although the existence of multiple binding conformations has been experimentally demonstrated(28, 34, 35), the two-states model is still controversial, as detailed in Ref. (31), where it was proposed that the speed-stability paradox is related to the use of continuum models for an intrinsically discrete problem.

Egr-1 (also known as zif268) is a TF responsible for the regulation of a variety of genes, and is induced by various stimulants, such as growth factors(36), neurotransmitters(37), hormones(38, 39), and stress(40). Previous studies have established a 9 bp ‘GCGTGGGCG’ consensus binding motif for Egr-1(41). A crystal structure of the protein bound to this motif shows that Egr-1 binds this recognition sequence as a monomer via interactions of its three zinc fingers (ZF1, ZF2 and ZF3), which contact 3 bp of DNA each(42). The wide interface created by the ZFs when bound to DNA, provide a strong, stable and specific binding. Upon stimulation, the levels of Egr-1 are estimated to rise up to ~10^4^ copies per nucleus(43, 44), but its lifetime is limited to ~1 h(44). Hence, in order to activate gene expression and respond to its stimulus, Egr-1 needs to rapidly scan the DNA in search for its response elements. During this search process, Egr-1 is required to discriminate its target site from among billions of bases, within a time frame of minutes. As a result, a tight binding conformation, like the one described above, can prevent the protein from efficiently scanning the genome, since many molecular bonds would need to be constantly created and disrupted. These seemingly contradictory requirements are a specific example of the ‘speed-stability’ paradox described above. Recent studies using NMR spectroscopy helped to resolve it, revealing a surprisingly different protein structure of Egr-1 while bound to non-specific DNA(45). In this conformation, ZF3 and ZF2 are bound to DNA, but ZF1 undergoes rapid dissociations. It was proposed that Egr-1 exploits this lower-affinity ‘scanning’ conformation in order to rapidly diffuse on the DNA, until it encounters its recognition sequence, binding to which traps the protein in the ‘recognition mode’ observed in the crystal structure. However, binding sites in real, transcribed genes do not exhibit in general the consensus sequence. In fact, many of these sites show gene-specific and evolutionary-conserved deviations from the consensus, indicating that the specific type of interactions Egr-1 makes with its binding elements have a functional importance. Hence, elucidating how the structure and dynamics of the Egr-1 complex is modulated by the binding site sequence and its genomic context is of great interest. Moreover, it is predicted that the mammalian genome harbors ~10^6^ sites that are highly similar, but not identical, to the classical recognition sequence of Egr-1(46, 47). Most of these sites, without a regulatory role, are expected to affect the search process, as they can momentarily trap Egr-1(46). The role of such sites is thought to be negative when these sequences are located relatively far from the target sequence, or positive if they are located in the vicinity of the response element where they help increase the protein’s local concentration(14, 15). Unfortunately, since most of the experiments have concentrated on the consensus sequence, it is not clear how Egr-1 binds to the variable ones. Interestingly, a thermodynamic additivity model was proposed to predict the affinity of near-consensus sites, but it was found to fail for sequences with four or more substitutions(47).

Two Egr-1 response elements reside ~60 bp apart within the evolutionary conserved proximal segment of the Luteinizing Hormone beta subunit (*Lhb*) gene promoter. In gonadotrope cells, Egr-1 is induced by the Gonadotropin Releasing Hormone (GnRH) to regulate the expression of the *Lhb* gene, and is essential for GnRH-induced gene activation and fertility in mice(48). Interestingly, both Egr-1 binding elements are different from the consensus motif, and they also differ by 4 bp from each other. These variations are conserved among the species (Supplementary Fig. 1), suggesting a functional role. Moreover, sequences flanking these sites are evolutionary conserved too, suggesting also that the DNA content surrounding these sites is important.

In this work, we used single-molecule DNA unzipping to study how the DNA sequence at the binding sites and their flanking regions alter the structure and affinity of the Egr-1 DNA complex. This approach provides a direct measurement of the position and forces associated with proteins bound to DNA(49–51), thus overcoming the inherent limitations of averaging in more traditional ensemble methods. We first measured the occupancy and breaking forces associated with Egr-1 binding to each of its response elements in their native context on the *Lhb* gene promoter, and then compared them with those obtained for the same sequences in other DNA contexts. We also used a novel method to characterize the dissociation of Egr-1 from its underlying DNA under asymmetric perturbations of the DNA. Together, the findings indicate that each of the functional binding sites, and their native flanking sequences, modulate the structure and affinity of the Egr-1-DNA complex in a different way, and suggest a unique functional role for ZF3 contacts with DNA.

## MATERIAL AND METHODS

### Reagents

The original plasmid for Egr-1 (DNA binding domain) was kindly provided by Dr. Scot Wolfe. The protein was expressed and purified as previously described(52), by cloning it into a pGex2t plasmid (GE Healthcare Life Sciences) along with a cleavable glutathione S-transferase (GST) tag. Fusion protein was expressed in BL21 cells grown in LB media at 37 °C, and purified using a glutathione sepharose column (GE Healthcare Life Sciences). GST tags were cleaved via room-temperature overnight incubation with thrombin (GE Healthcare Life Sciences) and then eluted. Purified protein (stock concentration 50 μM) was aliquoted and stored at -80°C until use. Specific binding of the purified protein to DNA was verified by EMSA with DNA oligomers containing the consensus binding site.

### Optical Tweezers

Experiments were performed in a custom-made double-trap optical tweezers apparatus(53), as previously described(8, 54). Briefly, the beam from an 852 nm laser (TA PRO, Toptica) was coupled into a polarization-maintaining single-mode optical fiber. The collimated beam out of the fiber, with a waist of w_0_=4mm, was split by a polarizing beam splitter (PBS) into two orthogonal polarizations, each directed into a mirror and combined again with a second BS. One of the mirrors is mounted on a nanometer scale mirror mount (Nano-MTA, Mad City Labs). A X2 telescope expands the beam, and also images the plane of the mirrors into the back focal plane of the focusing microscope objective (Nikon, Plan Apo VC 60X, NA/1.2). Two optical traps are formed at the objective’s focal plane, each by a different polarization, and with a typical stiffness of 0.3-0.5 pN/nm. The light is collected by a second, identical objective, the two polarizations separated by a BS, and imaged onto two Position Sensitive Detectors (First Sensor). The position of the beads relative to the center of the trap is determined by back focal plane interferometry(55). Calibration of the setup was done by analysis of the thermal fluctuations of the trapped beads(56), which were sampled at 100kHz.

### Molecular construct for single molecule experiments

The constructs for single-molecule experiments were generated as described previously(8), with a number of modification. Briefly, TF binding DNA segments with the *Lhb* promoter sequence were amplified by PCR from mouse genomic DNA, and segments for the non-native context (C1 and C2) were amplified from a plasmid containing the 601 nucleosome positioning sequence(57), a generous gift from Daniela Rhodes (MRC, Cambridge, UK). Primers used for the amplification reactions are listed in Supplementary Tables 8-9. For forward unzipping experiments, the constructs were digested using DraIII-HF (New England Biolabs) overnight according to the manufacturer’s instructions. A 10  bp hairpin (Sigma) was ligated to the construct using T4 DNA ligase (New England Biolabs), in a reaction with 1:10 molar excess of the hairpin, at 16 °C. The construct was subsequently digested overnight with BglI (New England Biolabs). For reverse unzipping the constructs were digested using BglI, hairpin-ligated and subsequently digested overnight with DraIII-HF. We generated two ~2000-bp DNA handles, each incorporating a specific tag (double digoxygenin and biotin), using commercially purchased 5’ modified primers in a standard PCR reaction, using bacteriophage lambda DNA as a template. The other two primers were designed to contain repeats of three DNA sequences recognized by single strand nicking enzymes: Nt.BbvCI and Nb.BbvCI (both from New England Biolabs) on the biotin-tagged handle and on the digoxygenin-tagged handle, respectively. The nicking enzymes generated 29 nt complementary overhangs on each handle. Handles were mixed at equal molar ratios for DNA annealing, creating a ~4000 bp fragment of annealed DNA handles. A ~350 bp dsDNA alignment segment with the sequence of the 601 DNA was prepared using commercially purchased primers (Supplementary Table 10) in a standard PCR reaction, ligated to the handles, and gel-purified (QIAquick 28706, Qiagen). Binding segments were ligated to DNA handles using a rapid ligase system (Promega) in 3:1 molar ratio, 30 min at room temp. The full construct (i.e. handles + alignement segment + TF binding segment) was incubated for 15 min on ice with 0.8 µm polystyrene beads (Spherotech), coated with anti-Digoxigenin (anti-DIG). The reaction was then diluted 1000-fold in binding buffer (10 mM Tris·Cl pH 7.4, 1 mM EDTA, 150 mM NaCl, 1.5 mM MgCl_2_, 1 mM DTT, 3% v/v glycerol and 0.01% BSA). Tether formation was performed *in situ* (inside the experimental chamber) by trapping an anti-DIG bead (bound by the DNA construct) in one trap, trapping a 0.9 µm streptavidin coated polystyrene beads in the second trap, and bringing the two beads into close proximity to allow binding of the biotin tag in the DNA to the streptavidin in the bead.

### Data analysis

Data were digitized at a sampling rate fs=2500 Hz, and saved to a disk. All further processing of the data was done with Matlab (Mathworks). Stretching of the tether was used to find the polymer-models parameters under our experimental conditions. Stretching to 15 pN was used to fit an extensible worm-like-chain model (XWLC) for the stretching of the dsDNA handles, and the data at forces above the unzipping of DNA was used to fit a worm-like-chain (WLC) model for the released ssDNA. From the measured and filtered tether extension and force, the stretching of the dsDNA handles at each time point was subtracted from the measured extension. Then, the extension was divided by the extension of two ssDNA bases (calculated from the measured force using the WLC model) to result in the number of unzipped bp. To improve the accuracy of the experiments, the alignment DNA segment was used to perform a correlation-based alignment of all traces in a group (i.e. a specific DNA sequence), allowing shifting of the traces (i.e. redefining the position of zero extension) and stretching of up to 2%.

### Measurements of breaking force and binding probability

The steerable trap was continuously moved to stretch the tethered construct to ~ 17 pN and unzip the DNA. In the presence of a bound protein, the propagation of the unzipping fork is halted and the force increases. At forces of 17 - 26 pN, the protein-DNA complex is disrupted in a single event that results in complete dissociation of the protein, leaving the DNA in a high-force, out-of-equilibrium state. A segment of DNA then immediately unzips, allowing the system to relax to equilibrium. Note, that the “width” of the peaks indicating bound proteins are thus related to the properties of DNA, but not to the size of the binding site.

After unzipping the whole construct, the DNA was relaxed to allow the re-formation of the dsDNA. This process was repeated multiple times, with a time between successive disruptions of a given site (the “incubation time”) of 40 s. The loading rate for the disruption of the proteins was 4 pN/s. Binding events were identified by detecting an increase in force of more than 0.5pN as compared to the median force at the same position, for experiments with the same DNA construct but in the absence of Egr-1. Applying the same criteria for the data obtained without Egr-1 resulted in no binding events detected. Events were classified as belonging to a specific site if the breaking event was located within a 20 bp window relative to the expected center of the binding site. Differences in breaking forces were checked using a two-tail Student’s t-test. Differences in binding probability were checked using a Chi-squared test. Differences were considered statistically significant if the calculated p-value was no larger than 0.05.

### Dissociation time measurements

DNA was unzipped until reaching an extension corresponding to the Egr-1 consensus binding site. The position of the unzipping fork was further adjusted to a mean extension and force that results in equal mean residency times between a locally ‘closed’ (dsDNA) and ‘open’ (ssDNA) configuration of the consensus DNA. The fluctuations in extension over time were measured for 1 min before the exposure to Egr-1, after which the construct was moved to a region of the laminar flow chamber (Lumicks) that contains Egr-1. Binding of the protein was detected by a sudden transition to a ‘closed’ state and a prolonged residence time in this configuration. No binding events were detected in the absence of protein (Supplementary Fig. 8a). The dissociation time was taken as the time lapsed until the reappearance of the fluctuations. Two different methods were used: In the first, following binding the construct was moved back to the channel in which Egr-1 was absent, to prevent re-binding, thus providing a single event of dissociation. In the second (Supplementary Fig. 8c) multiple binding and dissociation events were observed by keeping the construct at the Egr-1 channel.

## RESULTS

### Single-molecule probing of Egr-1 binding reveals distinct structural conformations

Two putative binding sites for Egr-1, with sequence CACCCCCAC and CGCCCCCAA, are located, respectively, at positions -41/-49 and -104/-112 on the mouse *Lhb* gene, and are both important for its induction in gonadotrope cells. These sites are remarkably conserved among the species despite a number of introduced substitutions which diverge from the consensus motif (Supplementary Fig. 1). To characterize the binding of Egr-1 to each of these sites in its native DNA context, we subjected the -517/+246 bp region of *Lhb* to single-molecule analysis by DNA unzipping. We attached each of the two DNA strands at the -517 end of the *Lhb* DNA to a ~ 2000-bp DNA ‘handle’ harboring a tag (Biotin and Digoxygenin, respectively), thus allowing to tether the DNA construct between two ~1 micro-meter microspheres (covered with streptavidin and anti-digoxygenin, respectively), trapped in a high-resolution, dual trap optical tweezers setup (Figs. 1a,b). Then, we subjected the construct to mechanical force by moving one of the traps away from the other. When the applied forces reached ~16-17 pN, the sample DNA unzipped, as evidenced by a reproducible pattern of events that include a sudden decrease in force together with an increase in extension. It has been previously shown(49–51) that following the propagation of the unzipping fork allows to measure the position and strength of protein-DNA interactions, as the force required to disrupt these interactions is significantly higher than those needed to disrupt DNA alone. Hence, using a laminar flow system, we exposed our construct to a solution containing the DNA binding domain of Egr-1 (referred to as Egr-1 for simplicity), and unzipped the DNA in its presence. As expected, a clear force elevation was observed in the positions proximal to ~-41/-49 (site -1) and -104/-112 (site -2), indicating binding of Egr-1 to these sites (Fig. 1c, Supplementary Fig. 2a). Surprisingly, an additional force peak was detected in the position surrounding ~-360 (site -3). When we analyzed the sequence of the DNA at this position, it was clear that it corresponds to an additional putative binding site located between -349/-358, which corresponds to GGCCCACTC, a motif with two base substitutions as compared to the 9 bp consensus, and different by 3-4 bp from the other Egr-1 binding sequences located on the *Lhb* promoter. The force peaks detected for the three sites were not observed in the absence of Egr-1 (Fig. 1d, Supplementary Fig. 2b), nor for a control DNA sequence that does not harbor any known Egr-1 binding sites. Moreover, although the experiment was performed at a high concentration of protein (500 nM), at which a significant amount of it will be bound to DNA non-specifically, the measured forces in non-specific regions were identical in the presence or absence of the TF, suggesting that the forces required to disrupt the non-specific complexes are significantly lower than those required to disrupt Egr-1 from the specific sites. In addition, to evaluate whether peaks detected in successive unzipping cycle for a single molecule correspond to new binding events following each unzipping cycle, we performed the following control experiment: We first unzipped the DNA, thus disrupting Egr-1 binding, and then moved into a flow channel that does not contain Egr-1, relaxed the tension to allow reannealing of double stranded DNA (dsDNA), and immediately unzipped the DNA again. No proteins were found bound to DNA in any of the binding sites in the second unzipping cycle of these experiments (n=10), indicating that the disruption of the complex is irreversible.

**Figure 1:**
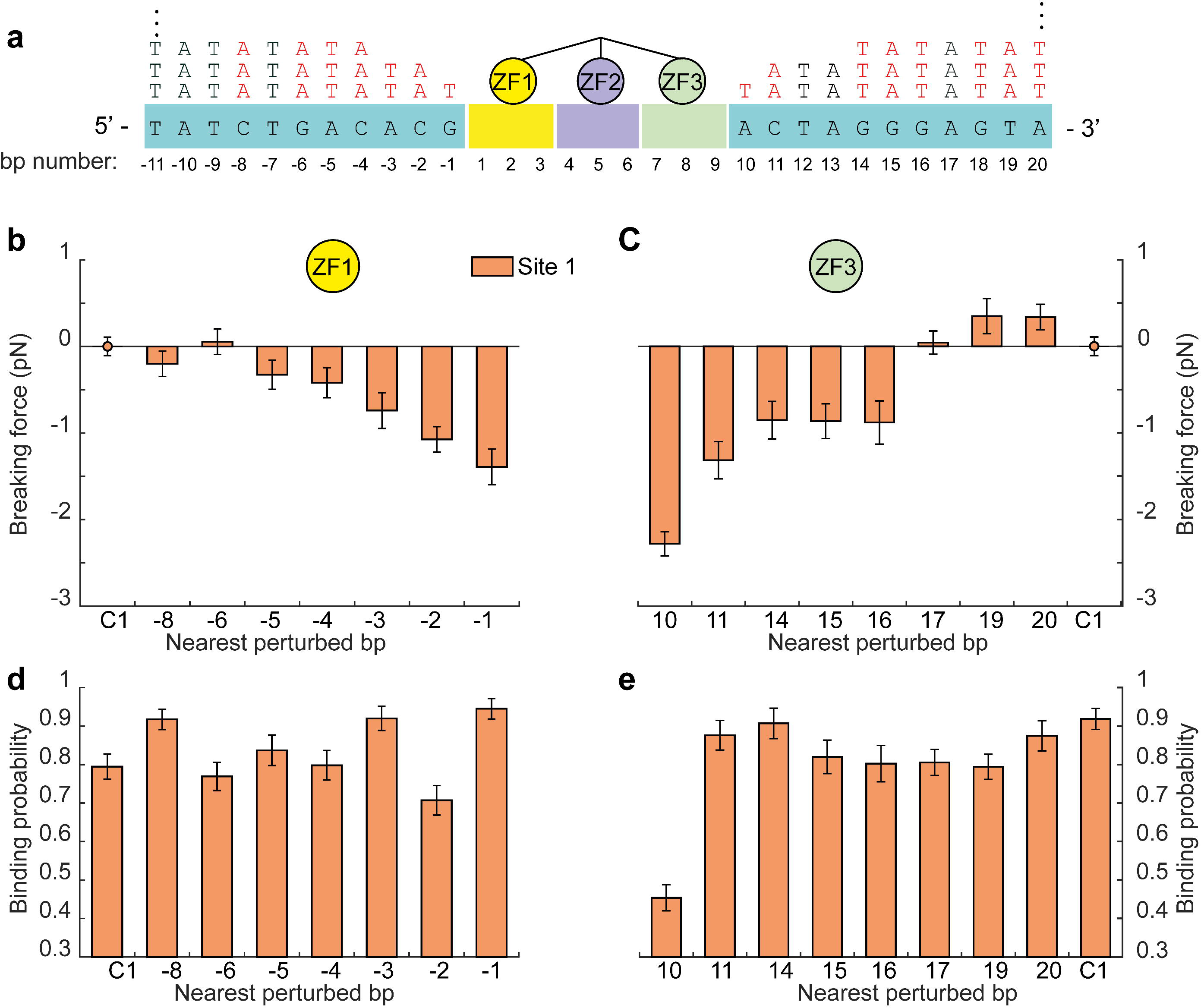
Single-molecule probing of Egr-1 binding to the *Lhb* promoter. (a) The -517/+246 segment of mouse *Lhb* DNA is ligated to a fixed 350 bp alignment sequence and a short stem-loop that prevents breaking of the tether after unzipping. (b) The *Lhb* DNA is connected to two 2kb dsDNA molecular handles, which are attached to polystyrene beads trapped in two separate optical traps. Egr-1 binding to the construct is initiated *in situ*(inside the experimental chamber). One of the traps is moved to stretch the tethered construct and disrupt protein-DNA interactions. After unzipping the whole construct, the DNA is relaxed and dsDNA forms again. The process is repeated multiple times. (c-d) Unzipping curves of *Lhb* in the presence (c) and absence (d) of Egr-1. Binding to each of the sites (-1/-2/-3) is designated with arrows.(e) Breaking force for the three Egr-1 binding sites located on the *Lhb* promoter. Data shown as mean ± s.e.m., *n*=140; ****p<0.0001, two-sample Student’s T-test. (f) Binding probability, calculated as the number of binding events out of the total number of DNA unzipping cycles, for each of the three Egr-1 binding sites at [Egr-1] = 500 nM. Data shown as fraction ± s.e., *n*= 140; ****p<0.0001, Chi-square test.

Accordingly, unzipping DNA multiple times in the presence of Egr-1, allowed us to calculate the mean breaking force (Fig. 1e) as well as the fraction of successive Egr-1 binding events, out of the total number of unzipping cycles (i.e. the binding probability; Fig. 1f), for each of the three sites. Remarkably, these experiments revealed significant differences in the binding probability, between each of the three sites: Egr-1 was found bound to site-1 with the highest probability, as compared to both -2 (p<0.001) and -3 (p<2x10-8) sites. Increasing the time between consecutive unzipping cycles did not affect the binding probability, indicating that the system is able to reach thermal equilibrium (Supplementary Fig. 9). Hence, the binding frequency measured in these experiments should reflect the affinity of Egr-1 for a particular DNA sequence(59). It was previously shown that the affinity of Egr-1 to site -1 is higher than that to site -2 and lower than the affinity of the Egr-1 consensus sequence(60), which is consistent with our results. Next, to further clarify the differences in affinity between the different sites, we repeated these experiments in the presence of various concentrations of Egr-1. It has been shown that the consensus sequence exhibits the highest affinity of Egr-1 to DNA(47). Therefore, in addition to *Lhb* DNA, we performed the same experiment with the consensus sequence, introduced into a DNA segment derived from the 601 nucleosome positioning (context C1, Supplementary Fig. 3). For all the four sites probed, the binding probability is consistent with a hyperbolic saturation curve, as function of protein concentration (Supplementary Fig. 4a), from which the affinity of Egr-1 for a specific site can be estimated. As expected, the affinity for the consensus sequence was the highest, as compared with the sites located on *Lhb*, with similar affinities for sites -1 and -3, and lowest for site -2.

Interestingly, we found that the mean breaking forces (Fig. 1e, Supplementary Fig. 2c-e) are not correlated with the binding probabilities: For example, although the binding probability for site -3 was significantly lower than that of site -1, the mean breaking force for site -3 was significantly higher (p<10^−28^). In addition, the mean breaking force did not show a concentration dependence (Supplementary Fig. 4b), suggesting that it is an intrinsic property of each site. Importantly, while the binding probability at equilibrium is expected to depend only on the protein’s binding energy, the magnitude of the breaking force measured in these unzipping experiments depends on the *shape* of the energy landscape for the mechanical disruption reaction, which depends on the structure of the protein-DNA complex. As a result, differences in breaking force can reflect differences in structure, even if the different structures have similar affinities. Hence, the breaking forces we observed for the three binding sites on *Lhb*, which are uncorrelated with their binding probability, suggest that Egr-1 binds each of these sequence motifs in a significantly different structural conformation.

Previous studies suggested that the presence of an additional binding site proximal to a given site, can reduce its binding probability, as each site will compete with the other for protein binding. Since site -1 and site -2 are positioned only ~60 bp away from each other, we checked whether binding to each of these sites affects binding to the other. To that end we calculated the conditional probability of detecting a protein at a given site, given that there is a protein at the second. When we compared unconditional with conditional binding probability of both sites, we did not observe any significant difference between them (Supplementary Fig. 5).

### Flanking sequences proximal to zinc finger 3 modulate Egr-1 binding

The observed breaking force and binding probability of each single site was measured in the context of others sites *in cis*. This made us wonder whether the observed differences are solely due to the identity (i.e. the sequence) of the core 9 bp sequence of each site, or also affected by the sequence context. To that end, we introduced the 9 bp motif corresponding to each of the three sites, separately, into the same flanking context of C1 (Supplementary Fig. 3), and subjected them to multiple rounds of unzipping in presence of Egr-1 (Fig. 2). These experiments revealed that, although the binding probability of site -1 was unchanged as compared to the one measured on *Lhb* (Fig. 2b), for site -2 it was significantly increased (p=4x10^−4^) and for site -3 significantly reduced (p=2x10^−3^). The breaking forces for site -1 and site -2 were significantly elevated, while for site -3 it was significantly reduced (Fig. 2a). Introducing the sites into a different flanking context (C2, Supplementary Fig. 3), showed a significantly different set of forces and binding probabilities (Fig. 2a,b). For example, the binding probability for site -3 in the C2 context is nearly equal to the one in the native *Lhb* context.

**Figure 2:**
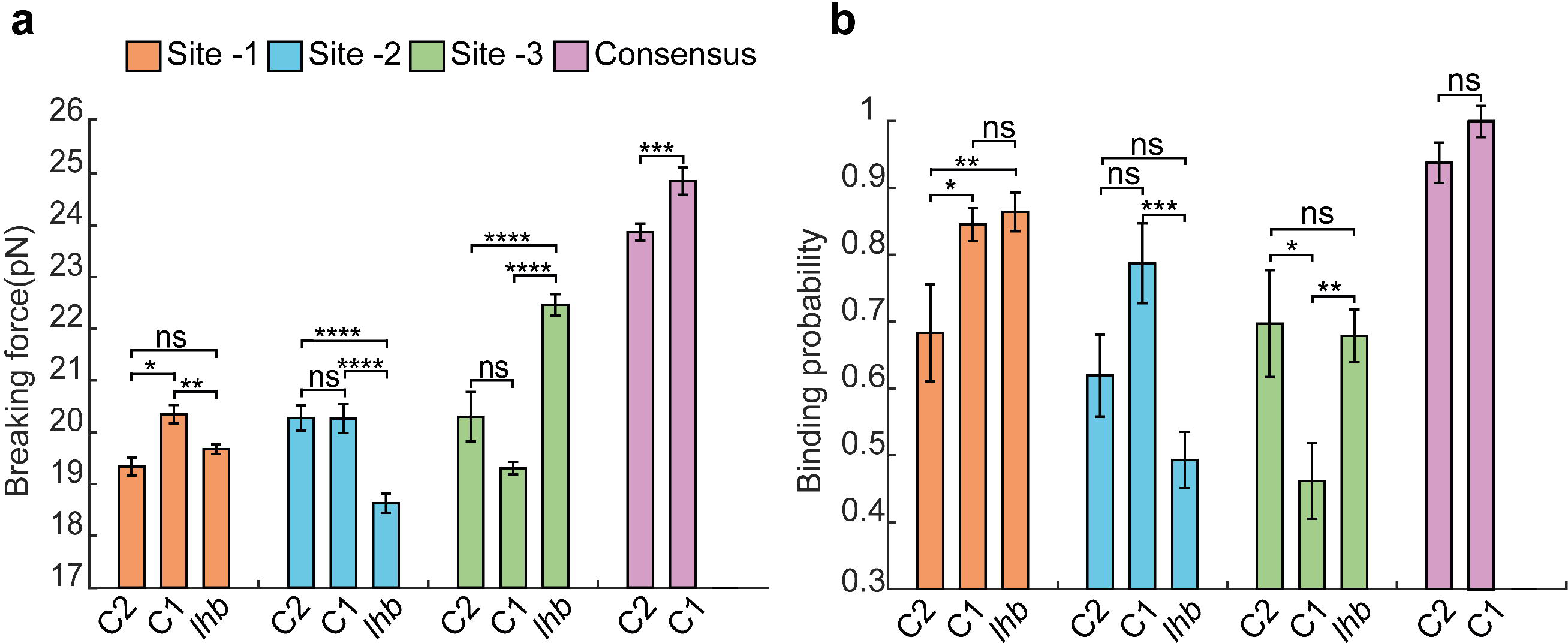
DNA sequence flanking the binding sites modulate Egr-1 binding. Three Egr-1 binding sites are integrated, separately, into a C1 and C2 DNA context (Supplementary Fig. 3) and subjected to multiple unzipping cycles in the presence of 500 nM Egr-1. (a) Mean breaking force and (b) binding probability are presented for sites -3, -2, -1 and the consensus motif. The data is analyzed as in Fig 1. The number of experiments for each case is shown in Supplementary Table 1, and the p-values in Supplementary Table 5.

In order to understand the effect of the flanking sequences on Egr-1 binding, we used site -1 in the C1 context and mutated the first base proximal to the nucleotide triplet bound by ZF1 (base number -1). The crystal structure of Egr-1 shows that ZF1 can make hydrogen bonds with this base(61), thus it is possible that the observed changes in binding probability and breaking force upon change of DNA context, are due to the substitution of this specific nucleotide. Remarkably, unzipping experiments using the mutated flanking base at this position led to a mild decrease in breaking force, but showed no significant change in binding probability (Figs. 3b,c). In contrast, when we mutated the first nucleotide proximal to ZF3 (base number 10), a significant reduction in both breaking force and binding probability was observed (Figs. 3b,c). Interestingly, when we performed the same experiments with the consensus sequence, we did not observe significant changes in binding frequency or breaking force upon mutagenesis of base number 10, and a reduction in force but no change in binding probability for mutagenesis of base number -1 (Figs. 3d,e).

**Figure 3:**
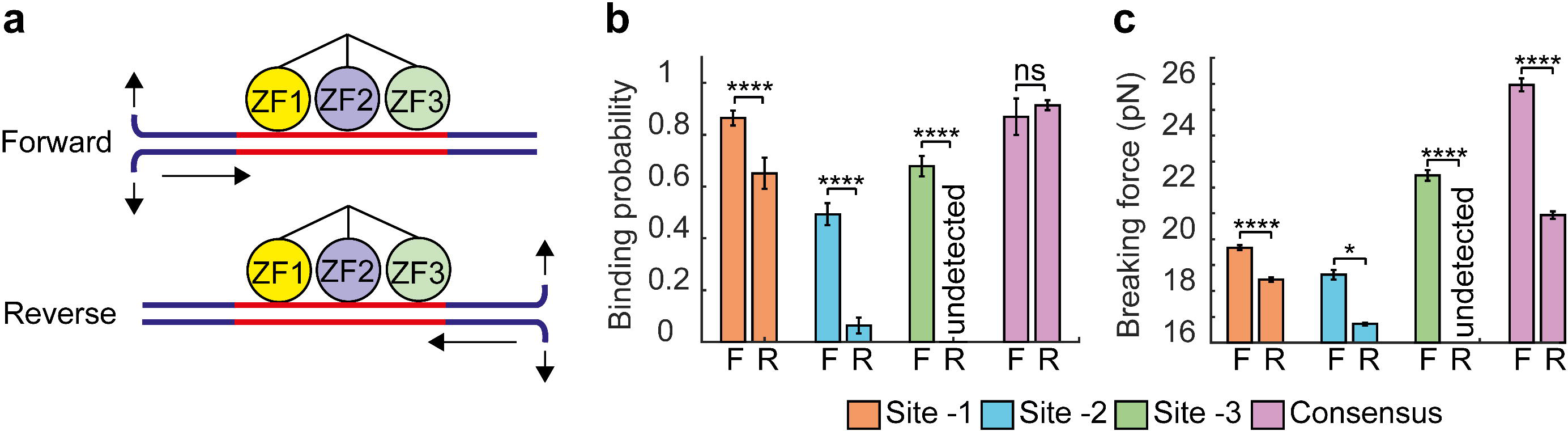
The first nucleotide proximal to ZF3 modulates Egr-1 binding. (a) The first nucleotide proximal to ZF3 or ZF1 is mutated to the indicated nucleotides. (b-e) Mutated DNA constructs for site -1 (b-c) or the consensus sequence (d-e), both in a C1 context, are subjected to multiple unzipping cycles in the presence of 500 nM. The data for binding probability and breaking force is analyzed and presented as in Fig 1. The number of experiments for each case is shown in Supplementary Table 2, and the p-values in Supplementary Table 6.

Previous studies have shown that 3-5 nucleotides surrounding the core binding site have a major effect on binding for some TFs(62, 63). In addition, a recent theoretical work predicted that the chemical composition near the binding site might influence the dynamics of search by a protein(64). Finally, genome wide data suggests that Zinc-finger TFs of the C_2_H_2_ type prefer GC rich over the AT rich flanking sequences(65). Hence, to check whether the effect of flanking sequences can extend beyond the first nucleotide flanking the binding motif, we gradually mutated 8-11 nucleotides proximal to ZF1 or ZF3 of site -1, by replacing them with repeats of the AT di-nucleotides (Supplementary Fig. 6). The effect of replacement on breaking force was evident for some of the nucleotide substitutions flanking both ZF1 and ZF3 (Fig 4). However, only the change of base number 10, proximal to ZF3, significantly reduced both breaking force and binding probability. Collectively, these results suggest that, at least for naturally occurring Egr-1 binding sites, the flanking sequences proximal to ZF1 affect the structure of the protein-DNA complex, without a strong effect on its affinity, whilst the first nucleotide proximal to ZF3 controls both structure and affinity of Egr-1-DNA complex. These results are consistent with a previous report(47), which measured the thermodynamic properties of binding to the consensus sequence containing single base-pair substitutions, and suggested the importance of ZF3 in the protein’s specific interactions with DNA.

**Figure 4:**
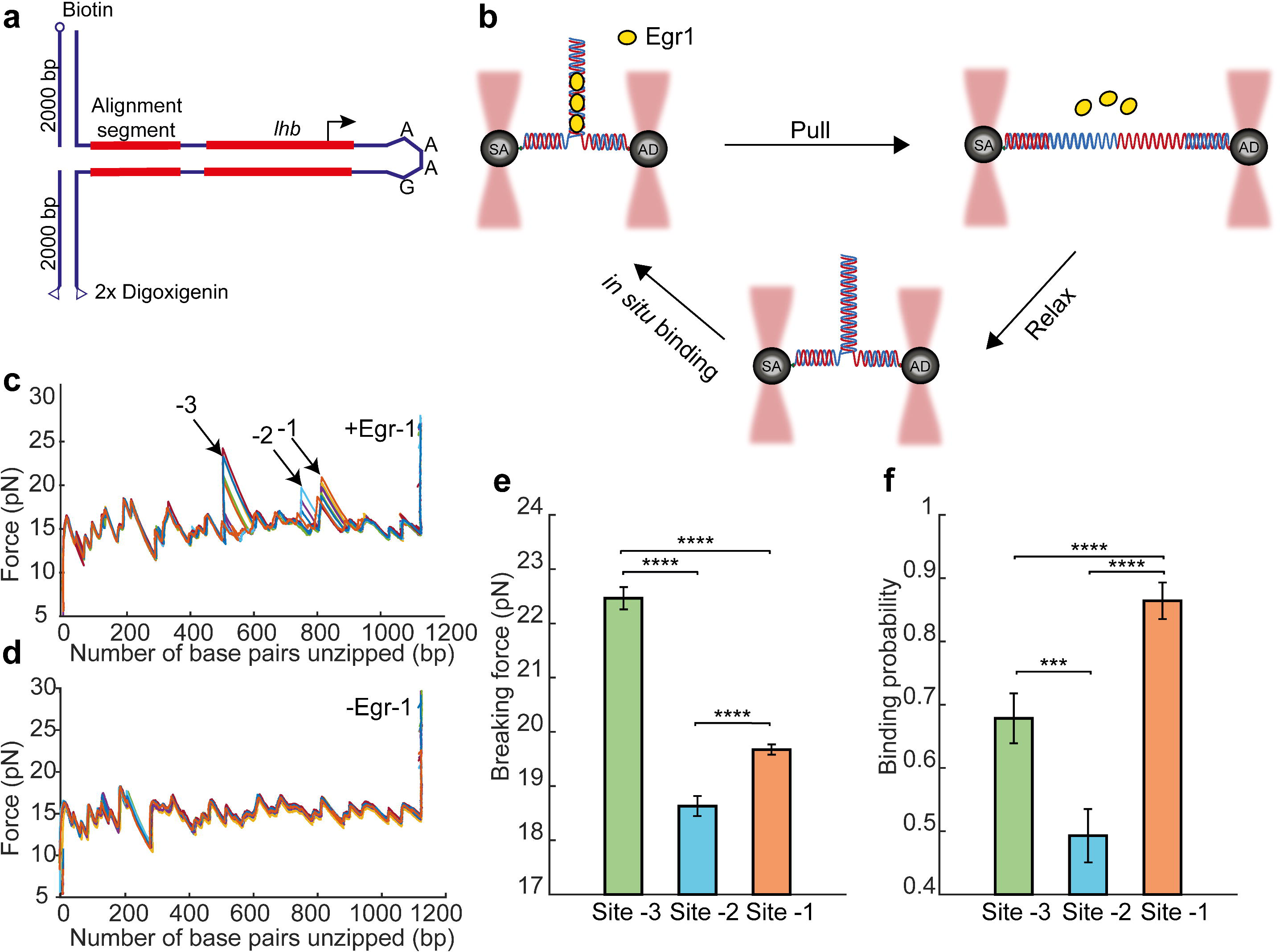
Effect for a gradual change of flanking context on Egr-1 binding. (a) The identity of 8-11 nucleotides proximal to ZF3 or ZF1, for site -1 in a C1 DNA context, is gradually changed by their conversion to AT repeats. (b-e) Mutated DNA constructs of site -1 are subjected to multiple unzipping cycles in the presence of 500 nM Egr-1. (b-c) Difference between the breaking force measured for the unmodified C1 context, and constructs in which nucleotides proximal to ZF1 (b) or ZF3 (c) were mutated. (d-e) The corresponding binding probabilities, analyzed and presented as in Fig 1. The number of experiments for each case is shown in Supplementary Table 3, and the p-values in Supplementary Table 7.

### Local disruption of ZF3 reduces Egr-1 binding

The observed asymmetric effect of flanking nucleotides suggested that the region proximal to ZF3 plays an important role in Egr-1 binding. Previous studies proposed that Egr-1 can bind to DNA via two distinct conformations, the recognition mode where all its ZFs are bound to DNA, and a scanning mode in which only two ZFs are bound (45, 66). Thus, we hypothesized that the dramatic effect observed as a result of the substitution of the nucleotide proximal to the triplet bound by ZF3, may reflect a particular sensitivity of the Egr-1-DNA complex for the local perturbation of ZF3, modulating Egr-1 binding as a whole. If this is the case, we would expect that dissociation of Egr-1 from the DNA will require less force if the perturbation is from the ZF3 direction.

To test this hypothesis, we compared the previous experiments unzipping *Lhb* from the -517 end (“forward unzipping”), where the unzipping fork encounters ZF1 first on sites -1, -2 and -3, with experiments where we unzipped *Lhb* DNA from the +246 end (“reverse unzipping”), approaching Egr-1 from the direction of ZF3 (Fig 5a). Notably, the mean breaking force required to disrupt the protein from sites -1 and -2 was significantly reduced (p<5X10^−11^ and p<0.02), as compared to the force required to disrupt it from the ZF1 orientation. Moreover, the force required for disruption of Egr-1 from site-3, that was the highest among the three sites in the forward unzipping, was so dramatically reduced, that we could not detect any force peak in the location near this site (Fig 5). This suggest that the sequence of the binding motif can modulate the specific interactions of the ZFs, creating different degrees of asymmetry in the structure for sites -2 and -3, where ZF1 is more tightly associated with DNA than ZF3, as compared to the complex formed on site -1 where the interactions are nearly symmetric. (Of note, the significant reductions we observe for the binding probability are likely the result of more binding events whose breaking force is close to the force sensitivity threshold set by the unzipping force of naked DNA, and are therefore missed). Surprisingly, two additional force peaks were detected in the positions previously unidentified inside the *Lhb* gene body (Supplementary Fig. 7). When we checked the identity of DNA sequence at the detected peaks, it was clear that they corresponded to the additional putative binding sites GAGTGGGTG and GAGGGGGTC at +113/+121 and +205/+213, respectively, downstream the TSS. Importantly, they are positioned in the opposite orientation as compared with sites -1,-2 and -3, so the unzipping fork first encounters ZF3 in these sites during “forward” unzipping.

**Figure 5:**
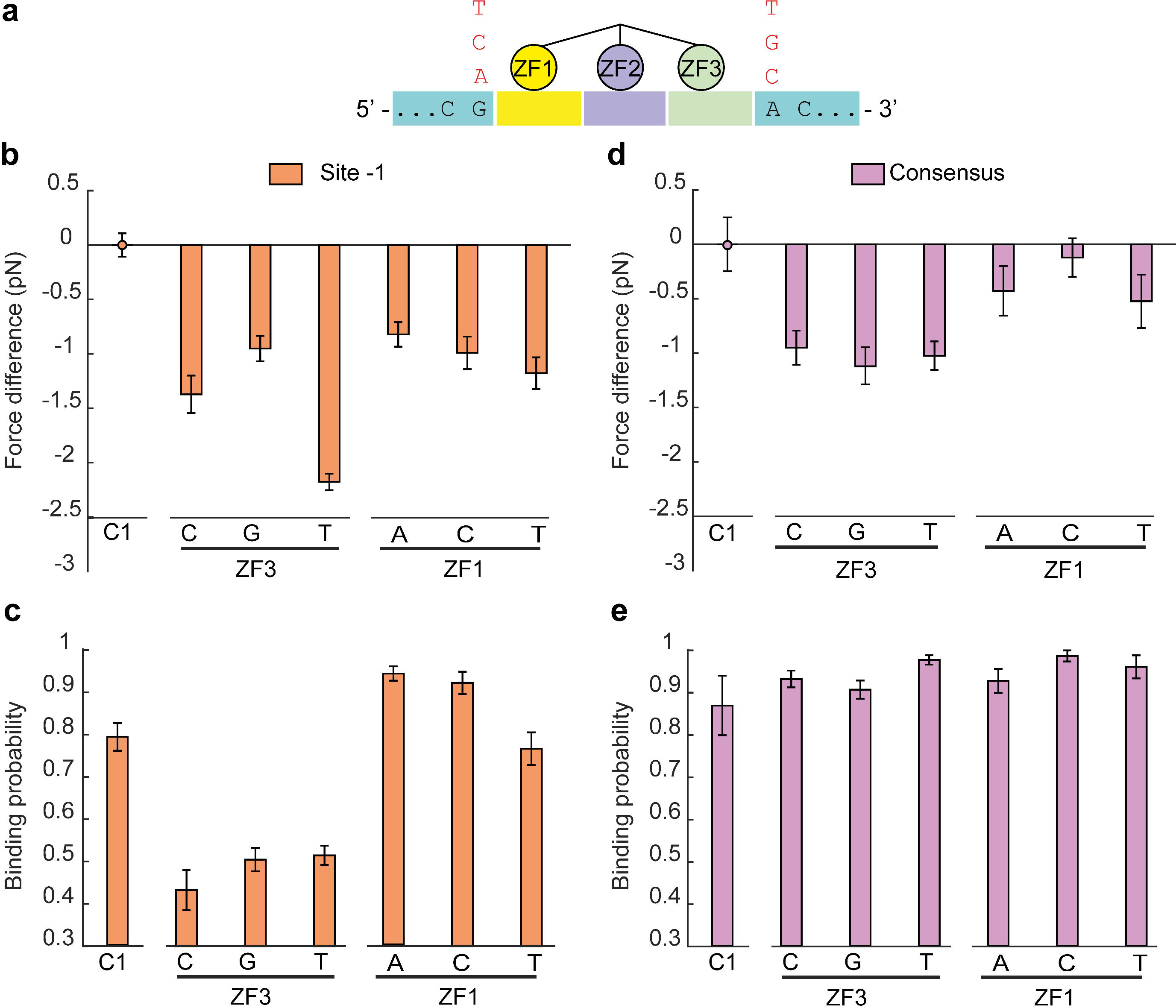
Unzipping Egr-1 from the ZF3 direction requires less force. (a) Schematic representation of forward and reverse unzipping. The *Lhb* promoter or the consensus motif in the C1 DNA context, are subjected to multiple cycles of forward (F) or reverse (R) unzipping in the presence of 500 nM Egr-1. The data for binding probability (b) and breaking force (c) is analyzed and presented as in Fig. 1. The number of experiments for each case is shown in Supplementary Table 1, and the p-values in Supplementary Table 5.

Next, to further check whether perturbation of Egr-1 from the ZF3 direction is a site-specific property, or a more general property of the structure of the Egr-1-DNA complex, we reverse-unzipped the Egr-1-consensus complex. Consistent with the results for the naturally occurring sites, we observed a significant reduction in breaking force (P<5x10^−22^) as compared to disruption from the ZF1 direction. These results suggest that the reduction in forces required for disruption of the protein through ZF3 is a general property that reflects Egr-1 binding to DNA.

Finally, we wanted to understand whether Egr-1 dissociation is faster upon a local disruption of ZF3, as compared with with a similar disruption of ZF1. Accordingly, we mimicked a local disruption of each ZF by partially unzipping DNA from the forward direction (for disruption of ZF1) or reverse direction (for disruption of ZF3). We unzipped the DNA until the fork reached the position of the consensus binding site, leaving it to rapidly fluctuate between a locally open (ssDNA) and locally closed (dsDNA) state (Fig. 6a,b). Next, we exploited our laminar flow cell to move the fluctuating DNA to the channel in which Egr-1 was present. Exposure of the DNA to the Egr-1 channel led to a quick transition of the DNA into a closed form, and a complete repression of the fluctuations, indicating Egr-1 binding. Reappearance of the fast fluctuations indicated dissociation of the protein. Multiple events of binding and dissociation could be observed for a single molecule with continuous exposure to Egr-1 (Supplementary Fig. 8c). To observe a single event of dissociation we moved the construct back to the channel in which Egr-1 was absent, to prevent re-binding. The time lapsed until the reappearance of fluctuations (Fig. 6b) allowed us to measure the characteristic dissociation time of a single Egr-1 molecule under a tension applied on the ZF1 or ZF3 side. Remarkably, when we approached the consensus from the ZF1 direction, the characteristic dissociation time was much longer (114 ± 3 s, mean ± sem) than the dissociation time measured when we approached the protein from the ZF3 direction (21 ± 1 s, mean ± sem; p=0.0004). Altogether, our results suggest that a local perturbation of ZF3, either by force or by modulation of the flanking sequences proximal to it, leads to more rapid dissociation of Egr-1 from its binding site.

**Figure 6:**
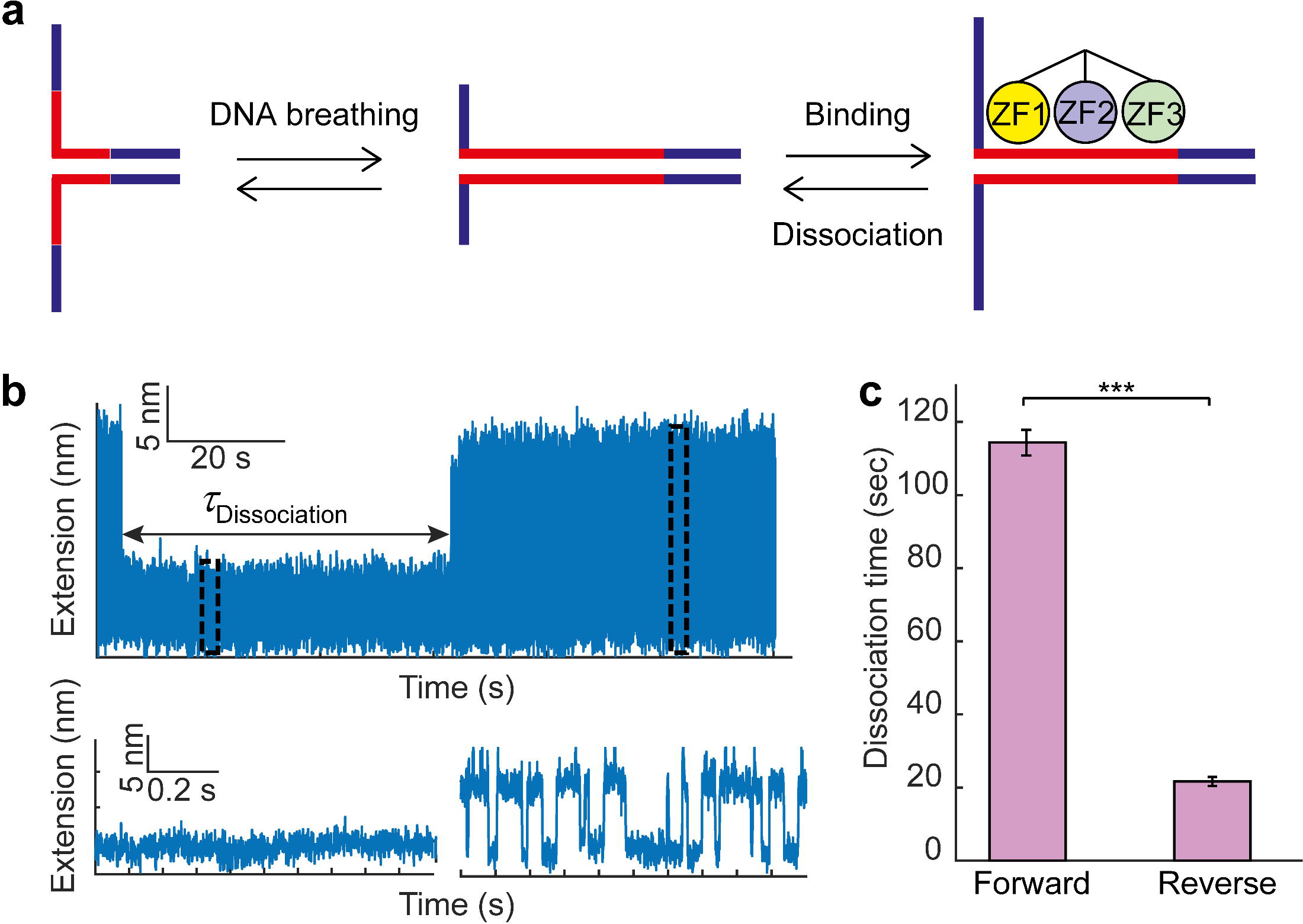
Force disruption of ZF3 increases Egr-1 dissociation. (a) A single-molecule containing a consensus motif (red) in the C1 context is unzipped until the fork reached the binding site. (b) The construct is held under tension, letting the DNA fluctuate between locally ‘open’ and ‘closed’ states.Binding of Egr-1 to the DNA stabilizes the closed conformation, leading to a sudden repression of the fluctuations, which lasts until Egr-1 dissociation. The time difference between binding and unbinding (dissociation time) is measured for forward (disruption of ZF1) and reverse (disruption of ZF3) unzipping. (c) Mean dissociation time, calculated for forward and reverse unzipping. Data shown as mean ± s.e.m., n = 25 and 14. ***p<0.001, two-sample Student’s T-test.

## DISCUSSION

Despite the universal requirement for gene-specific transcription factors to bind a highly specific recognition element in their target gene promoters in order to induce transcription, the identification and binding process is still surprisingly poorly understood. In this study, we have characterized the interactions of Egr-1 with its binding sites at the promoter of the *Lhb* gene. Measuring both the protein binding probability to the different sites and the mean force required to disrupt the bound proteins, allowed us to characterize the binding of Egr-1 at the single-molecule level. Our experiments show that the differences in sequence between these sites, and the specific genomic context where they are located, dictate different modes of interaction with the DNA. Interestingly, the breaking forces measured for the three conserved sites in the *Lhb* promoter did not correlate with their binding probability, suggesting that it is not only a modulation of the affinity of a site which is functionally important, but the specific structure of the complex, as dictated by the binding sequence and its context, is important too.

Our results show that the protein-DNA complex is particularly sensitive to the properties of the DNA flanking the binding site at the side corresponding to interactions made by ZF3. Unzipping the DNA in the reverse direction, thus approaching the complex from the ZF3 side, resulted in lower breaking forces, indicating that the complex is more easily displaced if ZF3 is disrupted first. A novel method monitoring binding of Egr-1 via the reversible repression of local fluctuations in the DNA allowed us to determine that the dissociation time of Egr-1 is much shorter under a perturbation on the ZF3 side than it is under a similar perturbation on the ZF1 side, further supporting a functional role for ZF3. Interestingly, recent studies have described two possible scanning-mode conformations(66): in the first, “scanning mode a”, ZF2 and ZF3 are bound to DNA while ZF1 is not. This is the conformation originally described in Ref. (45). In the second possibility, “scanning mode b”, ZF1 and ZF2 interact with the DNA, while ZF3 doesn’t. It is possible that the effect we observe here for the perturbation of ZF3, stems from the fact that a transition from “recognition mode” to “search mode” could be catalyzed by disrupting the interactions of either ZF1 or ZF3. Since we observe an increased sensitivity for a perturbation of ZF3, our results may suggest that “scanning mode b” is the preferred mode of scanning accessed from this binding sites.

Notably, our findings have implications for the interaction of Egr-1 with other TFs. Binding of a second TF to DNA in close proximity to the ZF3 side of a bound Egr-1 can facilitate its dissociation, thus effectively providing a mechanism of cooperativity between the TFs. Interestingly, site -2 in the *Lhb* gene is located immediately upstream of the binding site for Pitx1, and downstream from the binding site of SF-1. These three TFs have been found to act in a highly cooperatively manner in the GnRH-induced expression of *Lhb*(39, 67, 68), and the mechanisms outlined here may provide a molecular mechanism by which this cooperativity is achieved. Our results can also shed light on the interaction of Egr-1 with other cellular molecular motors translocating on DNA. The mammalian genome harbors ~10^6^ sites that are highly similar to the classical recognition sequence of Egr-1(46). Some of these sites are *bona fide* regulatory elements, but most will likely be located at the body of genes. Previous studies have highlighted the importance of these quasi-specific sites for the kinetics of the Egr-1 finding its regulatory targets, as they can slow down the search process(46). However, binding of Egr-1 at high density on the DNA will likely affect also the function of molecular motors, e.g. polymerases, that translocate on DNA, thus potentially affecting important processes such as replication and transcription. Interestingly, while previous works have shown that many helicases are capable of displacing proteins bound to DNA (see for example Refs. (69–71)), polymerases seem to be unable to perform this task. Our results suggest a mechanism that can assist in the displacement of the quasi-specifically bound Egr-1 proteins, provided that they are in the right orientation, such that the polymerase encounters the ZF3 side first. Interestingly, all the binding sites we detected on the gene body of *Lhb* (1,2,3 in Supplementary Fig. 7) are positioned in an orientation such as an RNA polymerase transcribing the *Lhb* gene will first encounter Egr-1 from the ZF3 direction.

Finally, our findings on Egr-1 shed light on novel elements likely affecting also the interaction of other TFs with the DNA. Previous studies analyzing genomic variation in humans(72) have shown that most significant genetic variants are located in non-coding regions, and that complex organismal phenotypes are the result of altered binding of TFs. However, only a small fraction of these variations in TF binding is caused by variants directly disrupting the TF recognition motif(73). Our current study indicates the potential role of even small changes in DNA sequence in regions flanking the previously recognized binding sites, and suggests that these could well form a broader mechanism involved in altering transcriptional regulation by DNA binding factors in general. Elucidation of these molecular mechanisms will lead to a clearer understanding on the organization of regulatory elements at gene promoters, will help understand inter-individual phenotypic variations and may improve our ability to dissect the molecular basis for disease susceptibility.

## ACKNOWLEDGEMENTS

We are thankful to Drs. Yael Mandel-Gutfreund, Tali Haran and Yaakov Levy for stimulating discussions.

## Author Contributions

S.R. and H.K. performed the experiments. O.M. and A.K. designed and built the optical tweezers setup. S.R., H.K. and A.K. analyzed the data. A.S. and A.M. prepared reagents. S.R., H.K. and A.K. wrote the paper. P.M. and A.K supervised the research.

